# Cross-attention and language models reveal the interpretability of functional predictions for the human olfactory receptor family

**DOI:** 10.64898/2026.08.10.744067

**Authors:** Yi-Feng Zhang, Zi-huai Xu, Chen-xi Gao, Su-Yang Duan, Gang Li, Chang Xu, Hui-Meng Lu

**Affiliations:** School of Life Science and technology, Northwestern Polytechnical University, Xi’an, China; School of Computer Science, University of Science and Technology of China, Hefei, China; College of Life Sciences, Shaanxi Normal University, Xi’an, China; Semiochemicals for Green Prevention and Control of Plant Diseases and Insect Pests, Ministry of Agriculture and Rural Affairs, Yangling, China

**Keywords:** attention, olfactory receptors, protein function prediction, key region

## Abstract

The attention mechanism offers a route to data-driven discovery of biological principles. However, for important protein families such as human olfactory receptors, the extent to which attention can associate with biologically meaningful key regions lacks systematic validation. In this study, using human olfactory receptors (ORs) as a model, we constructed CrossVOI, a VOC-OR interaction prediction framework based on protein language models and cross-attention. Based on this architecture, we systematically analyzed the attention distributions of CrossVOI and found that although site-level attention for individual ORs was not stably reproducible, the averaged attention consistently concentrated on the C-terminal region and non-conserved sequence regions. In addition, we quantified the allocation of attention on ORs and VOCs and, combined with ablation experiments, explored the effects of data representation and model architectural complexity on the OR–VOC interaction prediction task. The results showed that increasing representation and model complexity did not always improve model performance on the prediction task. In summary, we propose CrossVOI, an interpretable OR–VOC interaction prediction framework with predictive performance comparable to that of current state-of-the-art models, and analyze the interpretability of the attention mechanism for human ORs. This study provides insights into the interpretability of protein function prediction methods and may help clarify the biological relevance of attention, and may guide future experimental validation.

## Introduction

Odorant receptors (ORs), also referred to as olfactory receptors, serve as the molecular basis for organisms to perceive and recognize volatile organic compounds (VOCs) in the environment. At the molecular level, olfaction begins with the specific binding between ORs and VOCs. Different ORs can bind to multiple distinct VOCs, exhibiting a many-to-many matching relationship. Currently, only a limited number of OR-VOC interaction (VOI) relationships have been deciphered, such as insects^1–5^, humans^6–9^, and other mammals^10–12^. The functions of most ORs remain unknown. Furthermore, in-depth study of key regions in ORs also has practical significance. By focusing on key regions of ORs, we can design biomimetic peptides with specific recognition capabilities^13^ or rationally design OR-targeted drugs. Therefore, reliably predicting VOI relationships and, in the process, exploring key regions that influence function is an issue worth addressing.

For the problem of identifying key regions affecting VOI, it is nearly infeasible to perform functional experiments by mutating residues one by one due to the high time and economic costs. In recent years, great progress has been made in the structural study of ORs. Using cryo-electron microscopy, the structures of human OR–G protein complexes^14^ have been resolved. Some key residues have been identified. On the basis of resolved structures, the molecular mechanisms of VOI can be elucidated through molecular dynamics simulations^17, 18^, which also aids in the discovery of key OR regions. Nevertheless, time and cost remain insurmountable bottlenecks; for the vast majority of ORs, key regions remain unknown. Therefore, existing computational or experimental methods are insufficient to effectively identify key OR regions, and cannot provide information on functionally critical regions for OR engineering or protein design.

The attention mechanism was first proposed in the Transformer architecture^19^ and has now become an essential component of deep learning methods. Attention-based architectures have achieved success in various bioinformatics tasks, such as drug-target interaction prediction^20, 21^ and protein function annotation^22^. Beyond improving predictive performance, the attention mechanism also provides a degree of interpretability. The attention weights learned during model training correspond, to some extent, to the importance of input features ^23^ and may indicate region-level trends, providing hints about regions of interest. With the development of protein language models (PLMs), self-supervised training on ultra-large-scale protein datasets can learn deep semantic information of proteins, providing richer sequence representations^24, 25^. However, it has also been argued that self-supervised learning relying solely on protein sequences is insufficient to capture receptor-ligand interaction information^26–29^, and that wet-lab experimental approaches remain indispensable. Therefore, whether attention-based function prediction models can yield reproducible and interpretable attention trends at the region level for ORs, becomes a primary question that we need to address.

Here, we propose CrossVOI, a VOI prediction framework based on a protein language model and the cross-attention mechanism. We trained and tested the model on M2OR^30^, the largest available experimental human VOI dataset, and achieved predictive performance comparable to that of previous methods. Furthermore, we systematically analyzed the cross-attention distributions of ORs. The results showed that attention did not stably pinpoint ligand-binding interfaces or evolutionarily conserved sites; instead, the reproducible signal was an enrichment of averaged attention at the H8–C-terminal region and at non-conserved sites, a region-level trend learned from wet-lab response data. In summary, we have constructed an interpretable VOI prediction framework with performance comparable to current state-of-the-art models and systematically analyzed the distribution of attention scores, providing a reference for interpretability research and for understanding the role of representation and model complexity in protein function prediction. Additionally, CrossVOI may provide a reference for future experimental validation and mechanistic analysis of olfactory receptors.

## Results

### CrossVOI workflow

The workflow of the CrossVOI framework is depicted in Fig. 1a. For a given set of VOC-OR pairs, the amino acid sequences of ORs and the SMILES sequences of VOCs are respectively processed by a protein language model (ProtT5) pre-trained on large-scale datasets and a chemical language model (ChemBERTa-2) to generate feature representations for ORs and VOCs. Self-attention modules are then applied to extract the intrinsic feature information of ORs and VOCs. Following this, a cross-attention module is used to integrate the two sets of features, capturing the interaction information between them. Finally, the integrated feature vector is input into a multi-layer perceptron (MLP) model to predict the interaction outcomes of the given pairs.

**Fig. 1.**
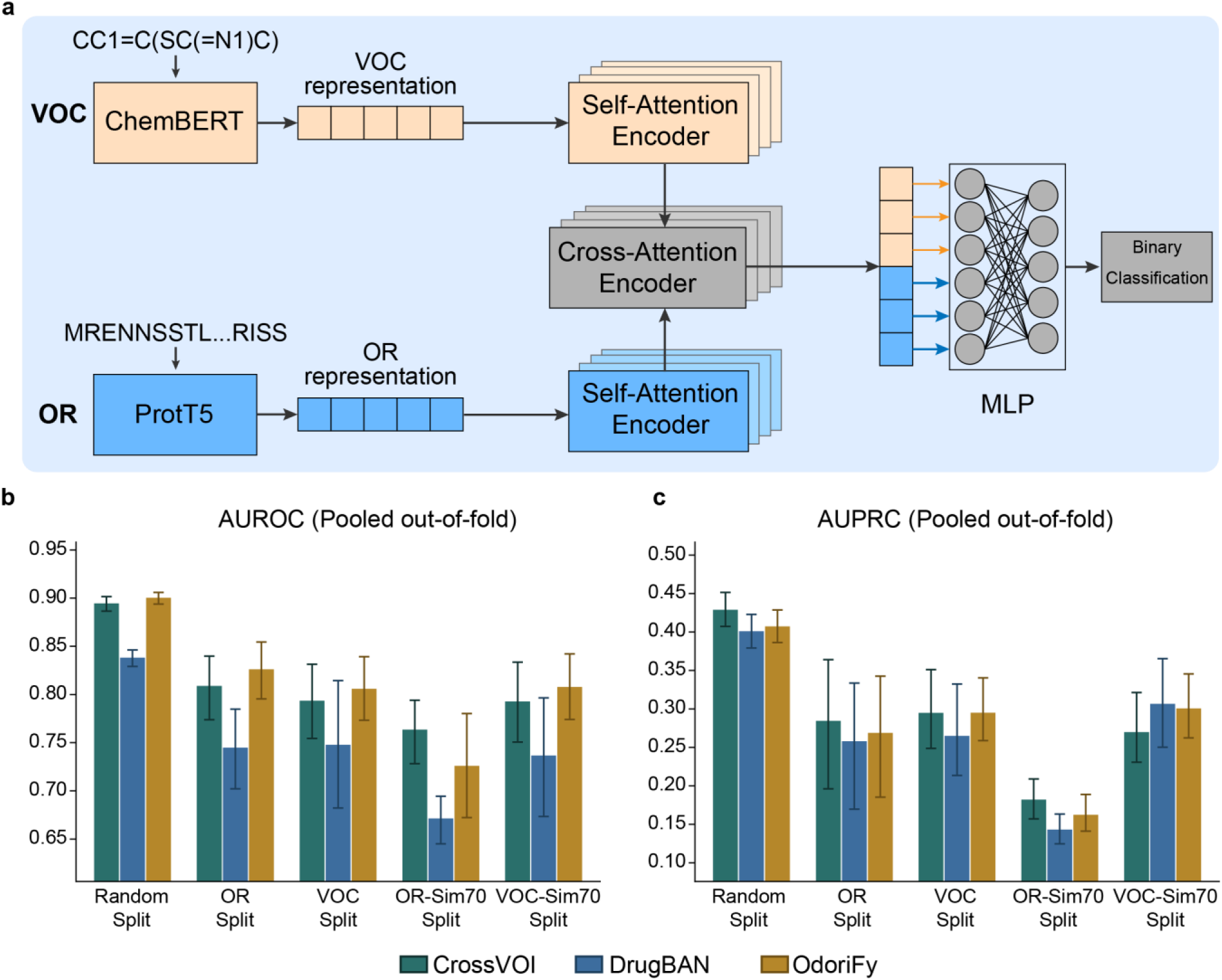
**a**: Architecture of CrossVOI. Sequence features from ORs and VOCs are encoded by language models, passed through self-attention and cross-attention, and then fed into a fully connected network. **b–c**: AUROC (b) and AUPRC (c) calculated from merged out-of-fold predictions across five folds; error bars represent 95% percentile intervals obtained by cluster-based resampling.

### Performance of CrossVOI

The experimental data were obtained from the M2OR database, which compiles experimentally measured responses for mammalian OR–VOC interactions. We used 42,917 human (Homo sapiens) entries from the processed Pair module (details in Materials and Methods) for model training and evaluation. For evaluating the binary classification task, we used AUROC and AUPRC as metrics.

As shown in Fig. 1b–c, under five different random seeds and five different data-splitting scenarios (random-split, OR-split, VOC-split, OR-sim70-split, and VOC-sim70-split; see Materials and Methods for details), we evaluated three frameworks by five-fold cross-validation—CrossVOI and two previously developed frameworks—including OdoriFy (a leading framework for human OR– VOC interaction prediction) and DrugBAN (a framework that performs well in drug–target interaction prediction). The results showed that CrossVOI and OdoriFy had comparable performance (OdoriFy had a slight advantage in AUROC, whereas CrossVOI had a slight advantage in AUPRC), while DrugBAN performed the worst, possibly because this framework was designed for drug–target interaction prediction and is not suitable for OR–VOC interaction prediction. We further discuss this point below.

### Overall distribution pattern of cross-attention across human ORs

We quantified the overall distribution of average cross-attention across OR sequences. As shown in Fig. 2a, cross-attention was unevenly distributed along the receptor sequence, with above-average attention scores in individual seeds at the N-terminus, C-terminus, and some transmembrane and loop regions. In particular, the C-terminus showed an above-average relative attention under different data-splitting scenarios; as shown in Fig. 2e, across different random seeds under multiple splits, the relative attention in this region was always greater than 1.

**Fig. 2.**
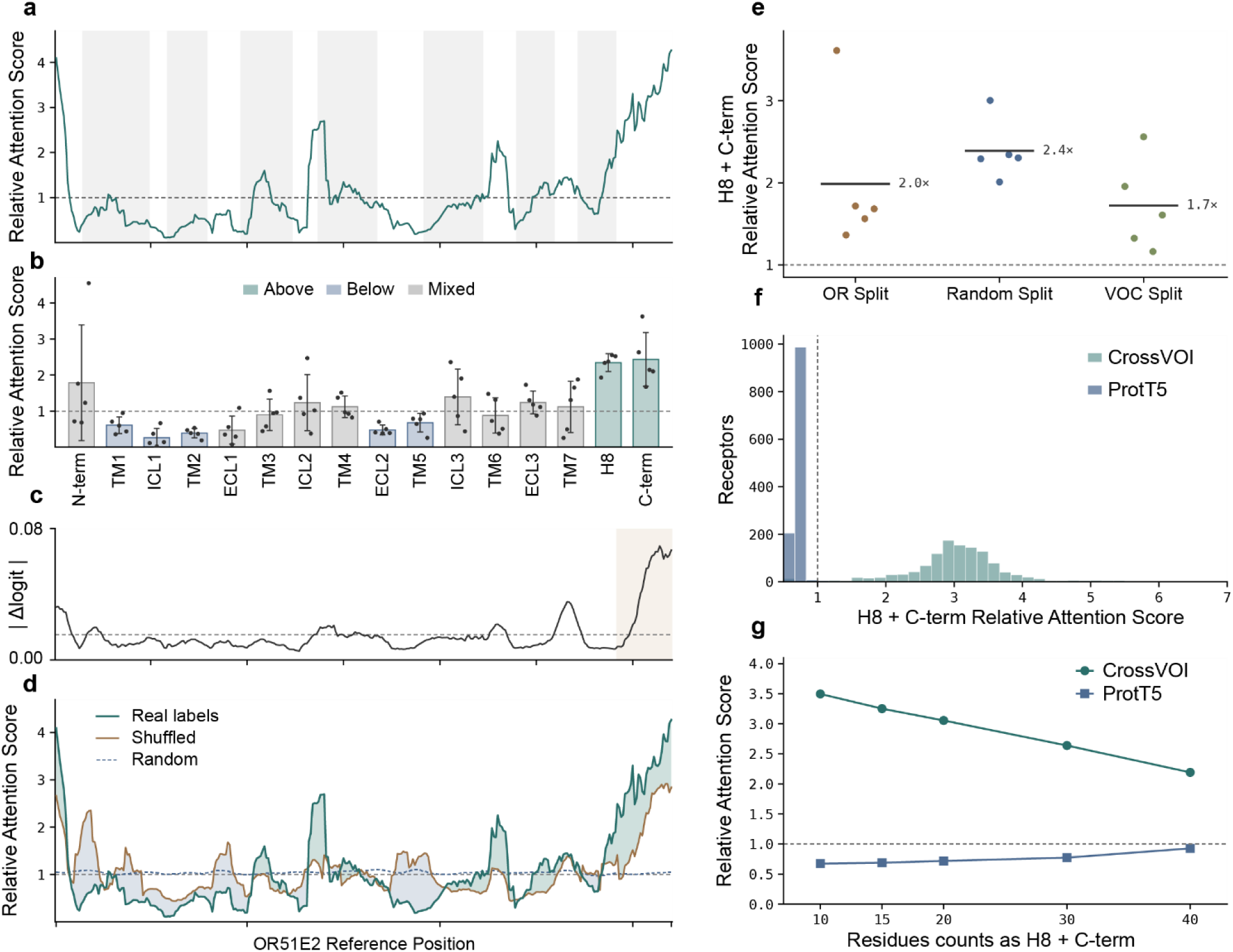
**a**: Residue-level average cross-attention along the OR51E2 reference axis. **b**: Average cross-attention for each structural domain; dots indicate values for the five training seeds, and colors indicate whether the direction was consistent across the five seeds (blue = all below 1, cyan = all above 1, gray = inconsistent). **c**: Absolute change in model logit after per-residue occlusion; the dashed line indicates the mean across the entire axis. **d**: Attention for three networks: trained with true labels, retrained with shuffled labels, and untrained. **e**: Average cross-attention on the H8–C-terminal segment for five random seeds under each split; horizontal lines indicate the mean. **f**: Attention proportion on the C-terminal 20 residues of 1,205 unique receptors, comparing trained cross-attention with frozen ProtT5 self-attention. **g**: CrossVOI cross-attention concentrates on the H8 + C-terminal region, whereas ProtT5 shows no attention to this segment; this conclusion is independent of sequence length.

Fig. 2b summarizes the segment-wise attention across five random seeds. Only the terminal H8 and the C-terminal portion showed above-average attention across all five random seeds, whereas TM1, ICL1, TM2, ECL2, and TM5 were all below 1; the direction of attention for the remaining segments was inconsistent across seeds. Therefore, only the C-terminus exhibited a stable and reproducible attention preference. To determine whether high-attention regions are important features affecting model output, we designed a simple occlusion experiment in which the features of the target region were replaced with the mean features over the entire sequence, and the change in model logit output before and after replacement, |Δlogit|, was calculated. As shown in Fig. 2c, the occlusion experiment demonstrated the importance of C-terminal features, as the mean |Δlogit| induced by occlusion within this segment was much higher than the average level.

To demonstrate that this preference arises from the data rather than the architecture, we examined the attention distributions of two control models, as shown in Fig. 2d. When the same architecture was randomly initialized but not trained, attention was nearly uniform along the sequence axis. After shuffling the training labels and fully retraining, the C-terminus still showed an elevated relative attention (up to approximately 3-fold the average level), indicating an intrinsic preference that does not require label information. Importantly, training with the true labels produced an even stronger C-terminal relative attention (up to approximately 4-fold the average level). This excess over the shuffled-label control shows that, beyond the intrinsic preference, a stable C-terminal relative attention is learned from the response data rather than being an inherent property of the architecture.

The model’s preference for the C-terminus may also arise from features encoded by the PLM. To rule out this confounding factor, we further compared the average cross-attention of CrossVOI with the self-attention extracted from ProtT5. The latter showed a median attention of only 0.72 on the C-terminal 20 residues of 1,205 receptors and was overall flat and slightly below uniform; in contrast, the median cross-attention of CrossVOI was 3.06, with 97.9% of receptors showing values greater than 1 (Fig. 2f). This conclusion was independent of C-terminal sequence length: when the number of C-terminal residues was increased from 10 to 40, the median cross-attention decreased from 3.50 to 2.19 but remained far above 1, whereas the self-attention of ProtT5 remained below 1 (Fig. 2g). Therefore, the concentration of attention toward the receptor terminus was independently learned by the model during training rather than arising from embeddings provided by the PLM.

### The averaged attention is associated with sequence conservation

At the single-receptor level, the cross-seed consistency of cross-attention was very low. As shown in Fig. 3b, when Pearson r was calculated for each of the 1,205 receptors between pairs of random seeds (averaged over the ten seed pairs), the median values for OR-split, Random-split, and VOC-split were only 0.07, 0.16, and 0.10, respectively. This indicates that the cross-attention scores of individual receptors are largely determined by the random seed and cannot be taken as evidence that they capture feature importance or biological patterns.

**Fig. 3.**
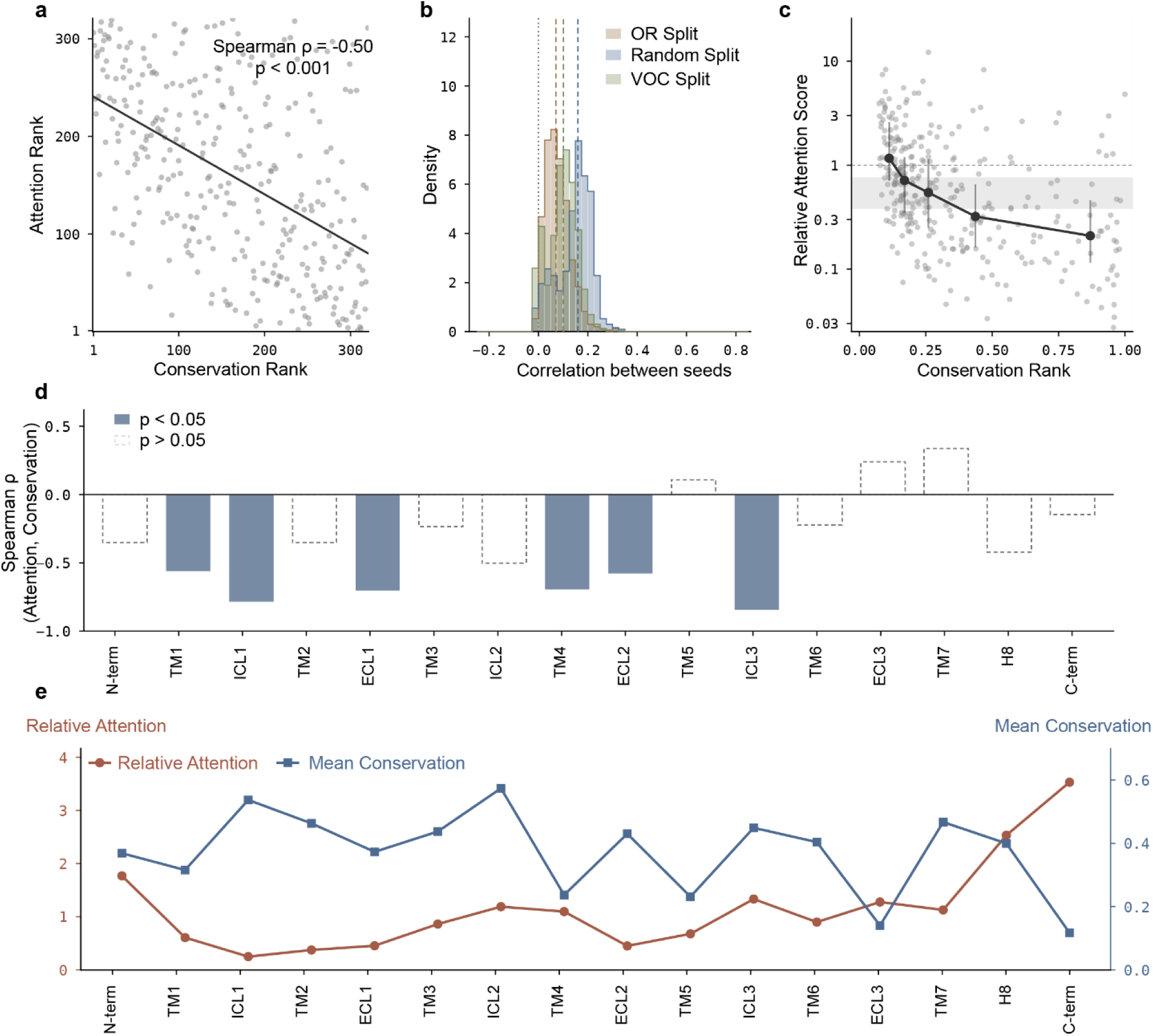
**a**: Spearman ρ = −0.50 between conservation rank (1 = most variable) and attention rank (1 = least attended) across 1,205 receptors. **b**: Pearson r of attention between two independent training runs (different seeds) for the same receptor; values were first averaged over seed pairs and then averaged over receptors, and dashed lines indicate the medians for each split (0.07, 0.16, and 0.10). **c**: Median relative attention (thick line) and interquartile range (vertical lines) for each quintile after binning by conservation; gray dots represent the raw values for 320 positions; the gray horizontal band represents the null hypothesis, i.e., the 95% interval of the quintile medians after randomly permuting the correspondence between attention and conservation 10,000 times. **d**: Spearman ρ between conservation and attention within each structural domain. **e**: Distribution of conservatism and attention

At the aggregate level, however, the average cross-attention displayed a degree of regularity. As shown in Fig. 3a, after aligning all OR sequences to the OR51E2 reference positions, attention was monotonically negatively correlated with conservation rank (Spearman ρ = −0.50, p < 0.001). When binned into quintiles by conservation, the relative attention decreased from 1.17-fold in the most variable quintile to 0.21-fold in the most conserved quintile; the medians of the quintiles from the random-permutation null distribution fell between 0.38 and 0.76, and both measured extremes fell outside this band (Fig. 3c). When calculated separately for the 16 structural segments, 13 segments showed negative correlations, 6 of which were significant; the three positively correlated segments were all non-significant and contained relatively few positions. The overall negative correlation was mainly contributed by segments such as TM1, ICL1, ECL1, TM4, ECL2, and ICL3. To some extent, these results suggest that attention focuses more on differences among input ORs than on their shared features.

### Greater feature and model complexity is not necessarily beneficial

In ablation experiments replacing the encoder, we found that the ligand encoder had almost no effect on model performance. As shown in Fig. 4b, the complex ChemBERTa-2 encoding and simple Morgan fingerprints did not yield a consistent improvement across splits, even under the ligand-targeted VOC Split and VOC-Sim70 Split. This may indicate that, for the current OR–VOC interaction prediction task, complex ligand feature representations do not provide additional useful information to the model, and simple chemical fingerprints are sufficient to provide ligand information for prediction. The attention similarity analysis in Fig. 4a further supports this conjecture. We compared attention similarity for the same OR paired with different VOCs and for different ORs paired with the same VOC. The results showed that replacing the OR markedly changed the attention, whereas replacing the VOC had little effect on attention, indicating that cross-attention does not focus on per-atom features in the ligand-side representation.

**Fig. 4.**
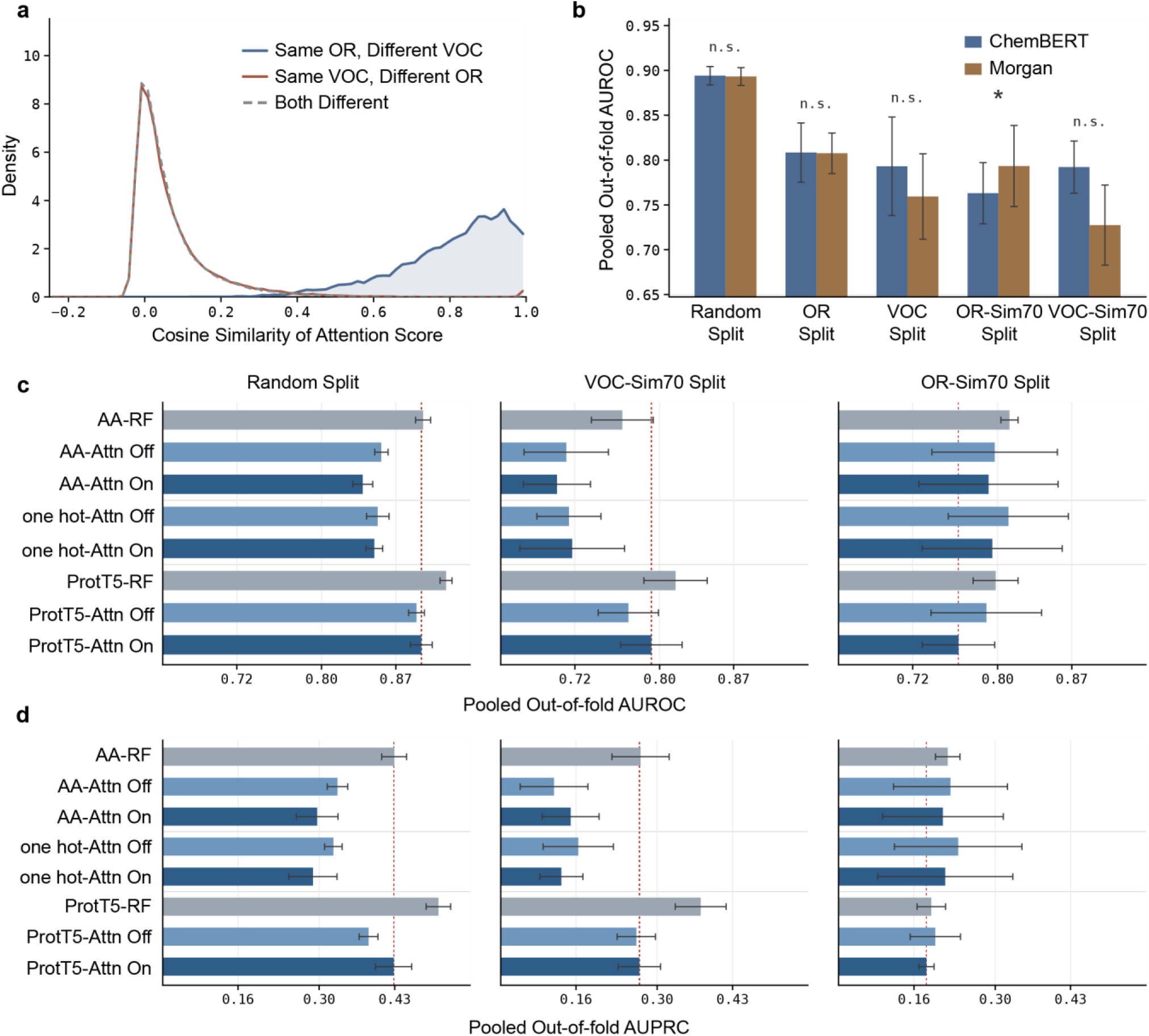
**a**: Cosine similarity of normalized cross-attention for each ligand at each layer. **b**: Performance comparison of models in which the ligand encoder was changed from ChemBERTa-2 to Morgan fingerprints. **c–d**: AUROC (c) and AUPRC (d) of out-of-fold predictions under three different data-splitting scenarios for combinations of protein-side representations and different model architectures.

Therefore, it is reasonable to speculate that, at least on the current dataset, there is an upper limit to the useful feature or model complexity for the OR–VOC interaction prediction task; that is, increasing the complexity of feature representations and model architectures does not always improve prediction accuracy. We present the performance of different protein-side representations and model architectures under different splitting scenarios in Fig. 4c–d. The results show that prediction performance does not increase monotonically with the complexity of representations and model architectures. The most complex combination of ProtT5 plus cross-attention (i.e., CrossVOI) never ranked first in the three splits. The combination of complex representations with a simple model (e.g., ProtT5 plus random forest) performed well across splitting scenarios, but the best combination depended on the metric and split (e.g., in VOC-Sim70 Split, ProtT5 plus random forest gave the highest AUPRC, whereas amino-acid composition plus random forest gave the highest AUROC); even in OR-Sim70 Split, which requires generalization to unseen ORs, this combination was the most stable. Moreover, the other combinations—simple representations with simple models, or complex representations with complex models—performed slightly worse. This suggests to some extent that features and models are not necessarily better when more complex; rather, there may be a reasonable range.

## Discussion

PLM performs self-supervised learning on ultra-large-scale protein data to mine semantic information from the protein world ^24^. However, it has also been argued that relying solely on the intrinsic information of proteins is insufficient to unravel the complex mechanisms of protein functional evolution, and that wet-lab approaches to reveal receptor-ligand interactions remain irreplaceable^26–29^. Our analysis supports this view to some extent. On the currently largest experimental dataset of human OR functions, however, the cross-attention of our model was consistently directed to the C-terminus and to sequence positions that are variable among ORs. Given that the C-terminus has been implicated in determining the response measured in OR functional assays, and that the model stably attended to this region, we hypothesize that this signal reflects information learned from the wet-laboratory response data. In other words, self-supervised learning on large-scale data alone cannot easily capture ligand-interaction information; therefore, ligand response data from wet-lab experiments are indispensable. A function prediction model based on cross-attention is expected to accelerate this process. This can be achieved, for example, by providing rapid preliminary screening for functional experiments or by mining potential key regions from experimental data.

Based on the distribution of attention scores, we examined from a different perspective the regions that may be relevant to human OR–VOC binding. However, it is worth noting that the attention mechanism is not a precise and reliable method for discovering key residues. For instance, although known key residues are often found in high-attention intervals, they do not always exhibit the highest attention scores. If the attention score were used as the sole criterion for identifying key residues, many false positives would arise. Therefore, the attention mechanism is better suited as a region-level indicator of importance trends rather than a residue-level precise locator. This limitation may arise from the size and quality of the dataset, as well as from inherent algorithmic constraints. The attention mechanism, originally designed for natural language processing (NLP), may not be fully compatible with the intrinsic patterns of protein sequences (e.g., protein function often involves the cooperative action of multiple residues, making the importance of individual residues difficult to isolate). Consequently, the attention mechanism is more appropriate for use in combination with other methods for identifying key residues, such as selection pressure analysis and molecular dynamics simulations, or for trend analysis at a broader scale, rather than for point-to-point precise localization.

In summary, we present CrossVOI, an interpretable OR–VOC interaction prediction framework with predictive performance comparable to that of current state-of-the-art models, and analyze the interpretability of its cross-attention for human ORs. Although site-level attention for individual receptors was not stably reproducible, the averaged attention was consistently concentrated on the C-terminal region and on non-conserved sequence regions. Quantifying how attention is allocated between ORs and VOCs, together with ablation experiments, further showed that increasing the complexity of representations or model architectures did not always improve prediction performance on this task. These results provide insights into the interpretability of protein function prediction methods and into how attention mechanisms relate to biological principles. We expect this framework to provide a reference for future experimental validation and to help clarify the biological relevance of attention in OR–VOC interactions.

## Materials and methods

### Dataset and data splits

The dataset was derived from the M2OR database^30^, which compiles experimentally measured mammalian OR–VOC interactions together with non-responsive experiments and experimental details. We extracted all functional-characterization results for Homo sapiens and binarized the outcomes into responsive and non-responsive according to the experimental system. Duplicate VOC–OR pairs were merged. All experiments used five-fold cross-validation with cluster-based folds, so that samples sharing the same receptor or ligand were not split across folds. Five splitting schemes were constructed to probe different generalization scenarios: Random-split (random assignment); OR-split (test receptors absent from training); VOC-split (test ligands absent from training); OR-sim70-split (holding out receptor clusters with no more than 70% sequence identity to training receptors); and VOC-sim70-split (holding out ligand clusters by similarity). The numbers of pairs, receptors and ligands in each split are provided in the Supplementary Materials.

### Model architecture and comparison protocol

The model was implemented in PyTorch^36^ (v1.12.0), with data processing in scikit-learn^37^ (v1.0.2), NumPy^38^ (v1.21.5) and pandas^39^ (v1.3.0). Maximum input lengths were set to 400 amino acids and 200 SMILES tokens, and shorter sequences were padded. OR sequences and VOC SMILES strings were encoded with ProtT5 and ChemBERTa-2, respectively; self-attention and cross-attention followed Monteiro et al.^20^, with implementation details given in the Supplementary Materials. For comparison, DrugBAN and OdoriFy were run using their official implementations without architectural modification. All three models used the same samples, the same fold assignments and the same input features, and were trained under an identical budget of at most 20 epochs with the best epoch selected by validation-fold F1; each model was trained once per split, so these are equal-budget rather than fully tuned comparisons.

### Evaluation metrics and statistical analysis

AUROC and the area under the precision–recall curve (AUPRC) were computed on pooled out-of-fold predictions: the five folds were concatenated and the metrics calculated once over all samples, rather than averaged across folds. Because only 5.4% of pairs are positive, AUPRC has no 0.5 chance level and was compared only within the same panel. Error bars are 95% percentile bootstrap intervals from 2,000 resamples at the cluster level (by receptor for OR-type splits, by ligand for VOC-type splits, and by sample for Random-split), which accounts for the correlation among samples sharing a receptor or ligand. Model comparisons used a paired cluster bootstrap on the same resampled clusters, and a difference was called significant when the 95% interval of the difference excluded zero. Since each model was trained once, these intervals describe test-set sampling error and not training-run variability.

### Attention extraction and reference axis

Cross-attention was extracted from out-of-fold predictions and averaged over the five folds and five seeds. Each receptor was aligned to the OR51E2 reference (L = 320 aligned positions) by global alignment with BLOSUM62, so that positions were comparable across receptors. Let *a_j_* denote the mean cross-attention at aligned position j after normalizing the attention profile to unit mass:

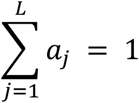

Hereafter, “relative attention” denotes the ratio of the observed attention at a position to that expected under a uniform distribution. The relative attention at position j was defined as:

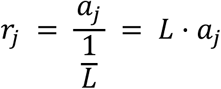

where the denominator is the per-position share expected under a perfectly uniform distribution. Thus a value of 1 indicates no positional preference, whereas values above or below 1 indicate enrichment or depletion relative to uniform. Sixteen structural segments (N-terminus, TM1–TM7, ICL1–3, ECL1–3, H8 and C-terminus) were defined from structural superposition and sequence annotation. A segment was called direction-consistent only when all five seeds fell on the same side of 1.

### Occlusion and control experiments

For 1,000 test samples, the feature at each residue was replaced by the receptor’s mean feature vector and the change in the model logit was recorded. Because occlusion pushed the logit up and down in roughly equal proportions, only the absolute change |Δlogit| was reported. Two controls used the same architecture, folds and training pipeline: an untrained network (forward pass only) and a network retrained after shuffling the training labels. To test whether the terminal preference was inherited from the protein language model, we extracted the last-layer self-attention of frozen ProtT5 for all 1,205 unique receptors, expressed it on the same relative-attention scale, and compared the fraction of attention in the terminal k residues for k = 10, 15, 20, 30 and 40.

### Cross-seed reproducibility and conservation

For each receptor, attention was represented as a mass profile over the aligned reference positions. Pearson r was computed between pairs of training seeds, averaged over the ten seed pairs and then over receptors, with profiles sharing fewer than 30 positions excluded. Profiles were decomposed into a per-seed group mean (shared shape) and receptor-specific deviations, and both correlations were reported. Sequence variability at aligned position j was quantified as:

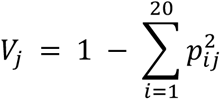

where *p_ij_* is the relative frequency of amino acid i at position j among the aligned receptors (i = 1…20), so that:

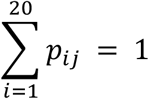

Conservation was defined as the complement of variability:

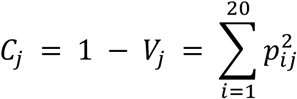

so that higher values indicate a more conserved position. Column statistics were estimated on rarefied samples of 600 receptors, repeated five times, to avoid low-coverage columns being dominated by few sequences. Attention and conservation were compared by Spearman’s rank correlation, quintiles were formed by equal-count binning, and significance was assessed against a null distribution obtained by permuting the attention–conservation pairing 10,000 times. Correlations within individual segments were interpreted by sign and p value only, as each contained 9–35 positions.

### Representation and encoder substitution

Protein-side representations (amino-acid composition, one-hot encoding and ProtT5 embeddings) were crossed with three model forms (random forest, a network without attention, and the full network) under a fixed ligand encoder, with each cell trained independently. The one-hot × random-forest cell was omitted because random forest requires a fixed-length vector and averaging one-hot would reduce it to amino-acid composition. In a separate control the ligand encoder was replaced by Morgan fingerprints with all other settings unchanged; because that arm has 14.9% more parameters, it is not a capacity-matched comparison. Detailed results and per-split sample counts are reported in the Supplementary Materials.

## Acknowledgements

This work was supported, in part, by the National Natural Science Foundation of China (32270525, 32470445); the Pherobio Semiochemical Institute Open Program (PHROBIO2023ZJSF02). We would like to thank all people in Lu Lab from Northwestern Polytechnical University for their approval.

## Author contributions

Y.Z., Z.X. and H.L. designed the research. C.G. and S.D. collected and analyzed literature data. C.G. performed the molecular docking. Y.Z. and H.L. wrote an initial draft of the manuscript. Further edits to the manuscript were provided by C.X. and G. L. All authors have given approval to the final version of the manuscript.

## Code availability

**Codes of CrossVOI download address:** GitHub (https://github.com/iORbase/CrossVOI).

