## supplement text for "Cross-attention and language models reveal the interpretability of functional predictions for the human olfactory receptor family"

**Data processing and dataset statistics**

The dataset was derived from the Pairs module of the M2OR database, which compiles experimentally measured mammalian OR–VOC interactions together with non-responsive experiments. The raw Pairs module contained 53,444 records. Dose–response measurements were preferred over screening data; contradictory dose–response results for the same pair were discarded (27 pairs). For screening data, pairs showing a response at a lower but not a higher concentration, as well as pairs with inconsistent results at the same concentration, were excluded (246 pairs); a consistent screening pair was labelled responsive if a response was observed at any tested concentration. After retaining only Homo sapiens records, 42,917 pairs remained, of which 2,319 were responsive (5.4%). All five splitting schemes used five-fold cross-validation with cluster-based folds. The numbers of pairs, receptors and ligands in each training/validation fold are summarized in Table S1.

Table S1. Number of samples in each fold under the five splitting schemes.

| Split | Fold | Subset | Rows | Positive Samples | Negative Samples | Unique Sequences | Unique SMILES |
| --- | --- | --- | --- | --- | --- | --- | --- |
| Random-split | 1 | train | 34333 | 1855 | 32478 | 1146 | 641 |
| Random-split | 1 | validation | 8584 | 464 | 8120 | 939 | 416 |
| Random-split | 2 | train | 34333 | 1855 | 32478 | 1150 | 624 |
| Random-split | 2 | validation | 8584 | 464 | 8120 | 936 | 428 |
| Random-split | 3 | train | 34334 | 1856 | 32478 | 1149 | 642 |
| Random-split | 3 | validation | 8583 | 463 | 8120 | 934 | 395 |
| Random-split | 4 | train | 34334 | 1855 | 32479 | 1167 | 624 |
| Random-split | 4 | validation | 8583 | 464 | 8119 | 920 | 418 |
| Random-split | 5 | train | 34334 | 1855 | 32479 | 1148 | 626 |
| Random-split | 5 | validation | 8583 | 464 | 8119 | 939 | 411 |
| OR-split | 1 | train | 33320 | 1684 | 31636 | 964 | 524 |
| OR-split | 1 | validation | 9597 | 635 | 8962 | 241 | 600 |
| OR-split | 2 | train | 34483 | 1851 | 32632 | 964 | 655 |
| OR-split | 2 | validation | 8434 | 468 | 7966 | 241 | 442 |
| OR-split | 3 | train | 34925 | 1970 | 32955 | 964 | 671 |
| OR-split | 3 | validation | 7992 | 349 | 7643 | 241 | 376 |
| OR-split | 4 | train | 34001 | 1934 | 32067 | 964 | 669 |
| OR-split | 4 | validation | 8916 | 385 | 8531 | 241 | 359 |
| OR-split | 5 | train | 34939 | 1837 | 33102 | 964 | 657 |
| OR-split | 5 | validation | 7978 | 482 | 7496 | 241 | 426 |
| VOC-split | 1 | train | 34717 | 2089 | 32628 | 1204 | 536 |
| VOC-split | 1 | validation | 8200 | 230 | 7970 | 886 | 135 |
| VOC-split | 2 | train | 34805 | 1827 | 32978 | 1188 | 537 |
| VOC-split | 2 | validation | 8112 | 492 | 7620 | 926 | 134 |
| VOC-split | 3 | train | 33649 | 1735 | 31914 | 1037 | 537 |
| VOC-split | 3 | validation | 9268 | 584 | 8684 | 1107 | 134 |
| VOC-split | 4 | train | 34550 | 1848 | 32702 | 1188 | 537 |
| VOC-split | 4 | validation | 8367 | 471 | 7896 | 880 | 134 |
| VOC-split | 5 | train | 33947 | 1777 | 32170 | 1148 | 537 |
| VOC-split | 5 | validation | 8970 | 542 | 8428 | 939 | 134 |
| OR-sim70-split | 1 | train | 34338 | 1856 | 32482 | 958 | 592 |
| OR-sim70-split | 1 | validation | 8579 | 463 | 8116 | 247 | 446 |
| OR-sim70-split | 2 | train | 34301 | 1860 | 32441 | 977 | 652 |
| OR-sim70-split | 2 | validation | 8616 | 459 | 8157 | 228 | 424 |
| OR-sim70-split | 3 | train | 34344 | 1853 | 32491 | 953 | 642 |
| OR-sim70-split | 3 | validation | 8573 | 466 | 8107 | 252 | 378 |
| OR-sim70-split | 4 | train | 34342 | 1853 | 32489 | 941 | 584 |
| OR-sim70-split | 4 | validation | 8575 | 466 | 8109 | 264 | 488 |
| OR-sim70-split | 5 | train | 34343 | 1854 | 32489 | 991 | 653 |
| OR-sim70-split | 5 | validation | 8574 | 465 | 8109 | 214 | 347 |
| VOC-sim70-split | 1 | train | 34388 | 1850 | 32538 | 1172 | 520 |
| VOC-sim70-split | 1 | validation | 8529 | 469 | 8060 | 889 | 151 |
| VOC-sim70-split | 2 | train | 34300 | 1891 | 32409 | 1055 | 522 |
| VOC-sim70-split | 2 | validation | 8617 | 428 | 8189 | 1043 | 149 |
| VOC-sim70-split | 3 | train | 34347 | 1841 | 32506 | 1198 | 536 |
| VOC-sim70-split | 3 | validation | 8570 | 478 | 8092 | 889 | 135 |
| VOC-sim70-split | 4 | train | 34378 | 1873 | 32505 | 1163 | 554 |
| VOC-sim70-split | 4 | validation | 8539 | 446 | 8093 | 932 | 117 |
| VOC-sim70-split | 5 | train | 34255 | 1821 | 32434 | 1142 | 552 |
| VOC-sim70-split | 5 | validation | 8662 | 498 | 8164 | 954 | 119 |

**Statistics of experimental systems in the original dataset**

We summarized the experimental systems used in the original dataset (Fig. S1). The assay systems were highly concentrated: Hana3A (H3A) cells with a luciferase (Luc) readout accounted for 58.6%, and human embryonic kidney (HEK) cells with a Luc readout accounted for 27.6%, together comprising 86.2%. When classified by cell line, co-transfection, and detection system, a total of 34 distinct protocols were observed. Among them, a single protocol using H3A cells stably expressing the receptor trafficking proteins RTP1/RTP2/REEP1 together with the Dual-Glo Luciferase assay accounted for 56.4%, whereas all other protocols were represented by only small numbers of samples.

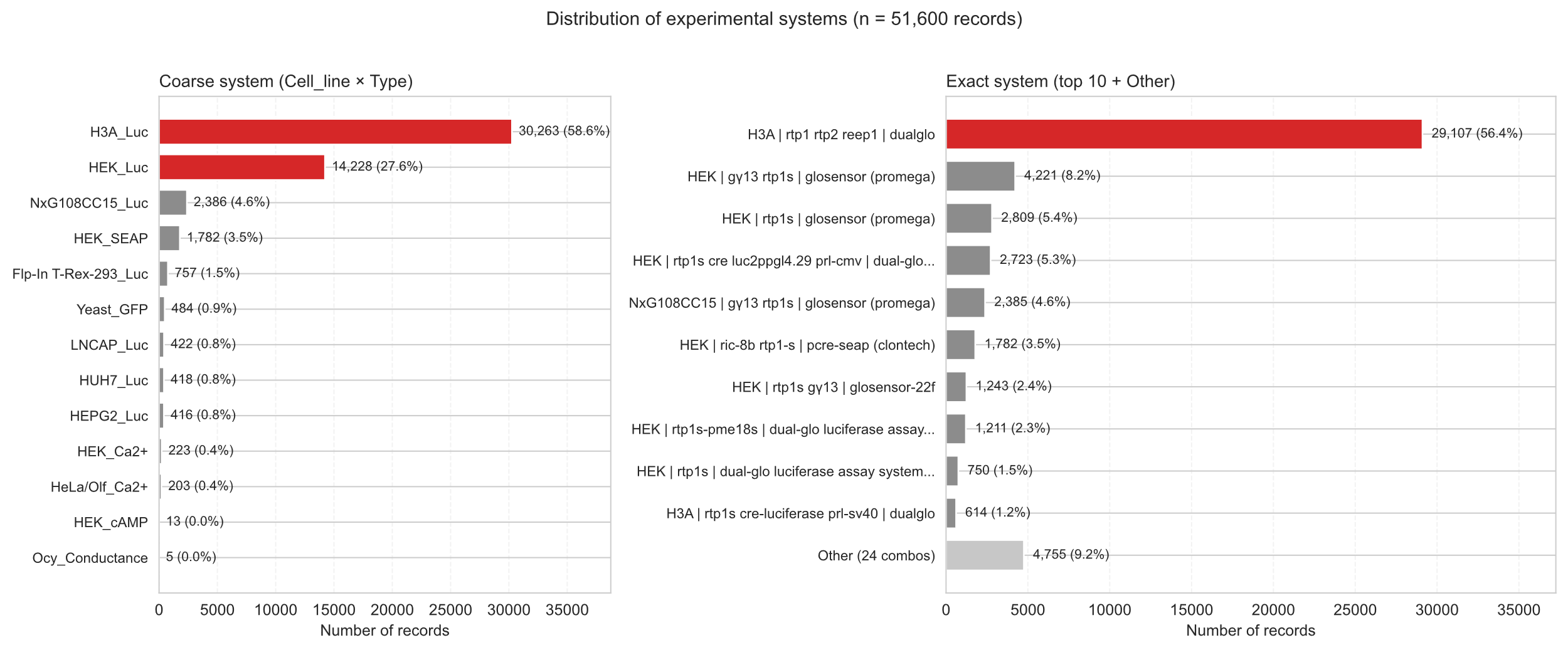

Fig. S1. Distribution of experimental systems in the original dataset.

**Molecular docking and contact frequency**

For each AlphaFold2-predicted OR structure, we aligned it to the cryo-electron microscopy structure of the OR51E2–G protein complex bound to a ligand. The docking box was centred on the ligand position of the cryo-electron microscopy structure and sized to fully cover the binding pocket. Docking was performed with AutoDock Vina v1.2.0. Because the binding pocket was defined in advance from the template structure, we docked the receptors against an independent collection of 2,686 VOCs reported in a previous large-scale study of olfactory functional evolution; this set was chosen because it covers a broader chemical space and is therefore more representative.

For each docked OR–VOC complex, residues within 5 Å of the ligand were counted; the contact frequency of a residue was the number of ligands placing it within this distance. Because the binding pocket was defined a priori in this docking workflow, the resulting contact frequencies approximate the spatial range accessible to ligands. As the contacts are derived from a computational model rather than experiment, they should be regarded as an estimate of the binding-pocket region and not as a strict definition of the pocket or as experimental validation of the attention profiles.

**Alignment of residues across different ORs**

To calculate the overall distribution of attention scores across residues of different ORs, we first performed multiple sequence alignment of the amino acid sequences of all human ORs. Using the GPCRdb domain annotation of OR51E2 as a reference, we mapped the attention scores of each OR to the corresponding domain positions of OR51E2 based on the alignment results. Specifically, for gap positions that exist in the OR51E2 sequence (i.e., positions where OR51E2 lacks a corresponding residue), the attention scores of other ORs at those positions were filled with zero. For gap positions present in the sequences of other ORs (i.e., residues that have no corresponding position in OR51E2), these positions were directly excluded from subsequent analysis. Through the above processing, we ensured that the attention scores of all ORs were aligned to the same domain reference for statistical analysis.

**Calculation of sequence conservation**

To quantify sequence variability at each aligned position, we used the definition introduced in the main text. After gap filtering, let the relative frequency of amino acid i at aligned position j be denoted as $p_{ij}$, where i indexes the 20 amino acids (i = 1…20), satisfying the normalization condition:

$$\sum_{i=1}^{20} p_{ij} = 1$$

Sequence variability and conservation at position j were then defined as:

$$V_{j} = 1 - \sum_{i=1}^{20} p_{ij}^{2}$$

$$C_{j} = 1 - V_{j} = \sum_{i=1}^{20} p_{ij}^{2}$$

Positions with more than 50% gaps were excluded. Column statistics were estimated on rarefied samples of 600 receptors, repeated five times, to avoid low-coverage columns being dominated by a few sequences.

**Details for attention computing**

The extracted feature vectors were then processed using self-attention and cross-attention modules to capture semantic information. The core attention computation is defined as:

$$Attn\left( Q,K,V \right)=softmax(\frac{QK^{T}}{\sqrt{d_{k}}})V$$

Compute the respective self-attention from the protein feature vector *P* and the small molecule feature vector *L* as follows:

$${SelfAttn}_{P}=Attn(PW_{Q},PW_{K},PW_{V})$$

$$P_{SelfAttn}=LN(P+{SelfAttn}_{P})$$

$${SelfAttn}_{L}=Attn(LW_{Q},LW_{K},LW_{V})$$

$$L_{SelfAttn}=LN(L+{SelfAttn}_{L})$$

The feature vectors processed by the self-attention module are then fused through the cross-attention module, where the cross-attention is computed as:

$${CrossAttn}_{PL}=Attn({SelfAttn}_{P}W_{Q},{SelfAttn}_{L}W_{K},{SelfAttn}_{L}W_{V})$$

$$P_{CrossAttn}=LN({SelfAttn}_{P}+{CrossAttn}_{PL})$$

$${CrossAttn}_{LP}=Attn({SelfAttn}_{L}W_{Q},{SelfAttn}_{P}W_{K},{SelfAttn}_{P}W_{V})$$

$$L_{CrossAttn}=LN({SelfAttn}_{L}+{CrossAttn}_{LP})$$

Here, ${CrossAttn}_{PL}$ denotes the cross-attention from protein to small molecule, and ${CrossAttn}_{LP}$ denotes the cross-attention from small molecule to protein. The key difference is that in self-attention, the queries (*Q*), keys (*K*), and values (*V*) are all derived from a single feature representation, whereas in cross-attention, the queries are derived from one feature representation while the keys and values are derived from the other.

**CrossVOI Hyper-parameter**

**Table S2. Model hyper-parameters.**

| Parameter | Value |
| --- | --- |
| Protein Encoder | ProtT5 |
| Ligand Encoder | ChemBERTa-2 |
| Protein Embedding Length | 400×1024 |
| Ligand Embedding Length | 200×768 |
| Self Attention Layer | 2 |
| Cross Attention Layer | 2 |
| Attention Head | 4 |
| Attention Dimension | 64 |
| Hidden Dimension | 256 |
| Dropout Rate | 0.1 |
| Learning Rate | 1×10⁻⁴ |
| Optimizer | Adam |
| Batch Size | 128 |
| Epoch | 20 |
| Random Seed | 1, 2, 3, 4, 5 |
| Loss Function | Binary cross-entropy |
| Early Stopping Metric | F1 Score |

**Key sites of OR51E2**

We categorized the key residues from previous work(Billesbølle, et al., 2023) into three functional classes: (1) ligand-binding pocket residues that directly contact the odorant; (2) G protein-coupling interface residues that mediate receptor–G protein interactions; and (3) activation switch residues that undergo conformational changes during receptor activation.

**Table S3. Key sites of OR51E2.**

| Type | Position |
| --- | --- |
| ligand-binding pocket residues | 262 |
|  | 258 |
|  | 181 |
|  | 104 |
|  | 155 |
|  | 158 |
|  | 198 |
|  | 202 |
|  | 180 |
| G protein-coupling interface residues | 121 |
|  | 217 |
|  | 243 |
|  | 291 |
| activation switch residues | 250 |
|  | 251 |
|  | 111 |
|  | 150 |
|  | 209 |

To test whether cross-attention was specifically enriched at residues with experimentally characterized functions in OR51E2, we repeated the analysis after completely excluding OR51E2 from the dataset. Using the five models from five-fold cross-validation, we quantified, for each residue, the fold change of cross-attention relative to the sequence-wide average. Almost none of the residues showed an attention fold change consistently greater than one (Fig. S2), indicating that the apparent enrichment at the previously reported key sites was not reproducible and could not be distinguished from random expectation. Accordingly, we removed this unstable result from the main text and discuss the interpretability of residue-level attention more cautiously.

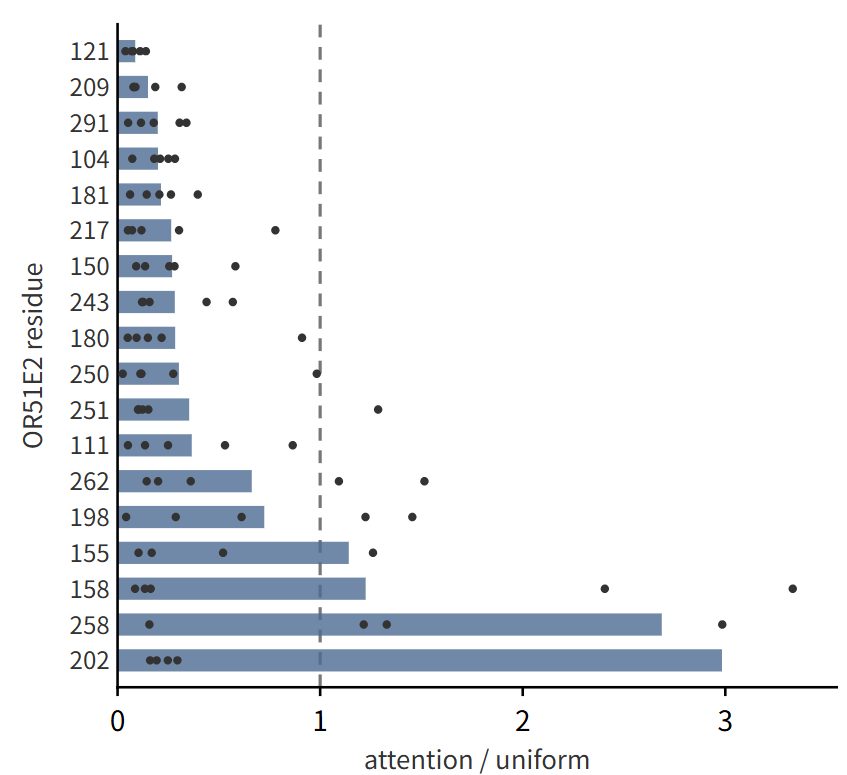

Fig. S2. Cross-attention at the 18 experimentally characterized OR51E2 residues. The dashed line denotes unity, i.e. attention equal to the sequence-wide average. Few residues show a fold change consistently above one across models, indicating no significant enrichment at the known key sites.

**Comparison with previous methods**

All three models (CrossVOI, OdoriFy and DrugBAN) were run with their official implementations without architectural modification. They used the same samples, the same fold assignments and the same input features, and were trained under an identical budget of at most 20 epochs with the best epoch selected by validation-fold F1. Each model was trained once per split.

Independent test set. As an additional external evaluation, we used a published dataset comprising 60 measurements of 3 ORs and 20 VOCs, of which 28 pairs were absent from the training data. The five models from the five folds of the Random-split were averaged to produce predictions on this test set. The model achieved AUROC = 0.797 and AUPRC = 0.802 on all 60 measurements, and AUROC = 0.711 and AUPRC = 0.664 on the 28 unseen pairs. Because the positive rate of this test set is 50%, the chance level of AUPRC is 0.5. The raw data are provided in the Independent_Test_Set sheet of Supplementary_Data.xlsx. This test set is limited in size and novelty, and a more comprehensive independent evaluation remains necessary.

Supplementary_Data.xlsx accompanying this document contains the underlying tables:

Orgin_Exp_Data (processed experimental records)

Sample_Partitions (fold assignments)

Baseline_Compare_Metrics (CrossVOI, OdoriFy and DrugBAN)

Per_Fold_Statistic (per-split sample counts)

Independent_Test_Set

Domain_Attention

Residue_Attention

Attention_VS_Conservation

Ligand_Encoder_Swap

Complexity_Combos.

Bai, P.*, et al.* Interpretable bilinear attention network with domain adaptation improves drug–target prediction. *Nature Machine Intelligence* 2023;5(2):126-136.

Billesbølle, C.B.*, et al.* Structural basis of odorant recognition by a human odorant receptor. *Nature* 2023;615(7953):742-749.

Chi, H.*, et al.* Genomic and phenotypic evidence support visual and olfactory shifts in primate evolution. *Nature Ecology & Evolution* 2025.

Eberhardt, J.*, et al.* AutoDock Vina 1.2.0: New Docking Methods, Expanded Force Field, and Python Bindings. *Journal of Chemical Information and Modeling* 2021;61(8):3891-3898.

Gupta, R.*, et al.* OdoriFy: A conglomerate of artificial intelligence–driven prediction engines for olfactory decoding. *Journal of Biological Chemistry* 2021;297(2).
